# Obsessive-compulsive disorder and abstract sequence task contributions shift prefrontal cortical connectivity

**DOI:** 10.64898/2026.05.22.727273

**Authors:** Hannah Hyde, Nicole C.R. McLaughlin, Sarah L. Garnaat, Theresa M. Desrochers

## Abstract

Obsessive-compulsive disorder (OCD) is characterized, in part, by repetitive, sequential behaviors, such as cleaning rituals, yet underlying neural circuitry related to abstract sequencing in OCD remains poorly understood. Prior work has implicated a set of cortical regions activated during abstract sequences, which are defined by rules rather than specific stimulus features (Desrochers et al., 2022). These regions include the rostrolateral prefrontal cortex (RLPFC) that is necessary for performance on abstract sequence tasks (Desrochers et al., 2015), as well as the anterior cingulate cortex/dorsolateral prefrontal cortex (ACC/DLPFC), supplementary motor area (SMA), middle temporal gyrus (MTG), and temporo-occipital junction (TOJ) that are differentially activated in OCD compared to healthy participants during abstract sequencing (Doyle et al., 2026). It remains unclear, however, whether these regions form a coordinated circuit, and how their interactions may differ in OCD. In the present study, we examined task based functional and effective connectivity among these regions using a previously published dataset. We tested hypotheses that connectivity within this circuit would be altered in OCD relative to healthy controls (HCs), and that prefrontal regions (ACC/DLPFC and RLPFC) would direct information to downstream regions (SMA, MTG, and TOJ) during a sequential task. We found that connectivity within this circuit differed significantly between groups. HCs exhibited less negative connectivity from the ACC/DLPFC to the TOJ and stronger positive coupling between the MTG and TOJ, as well as stronger coordination between the RLPFC and DLPFC, suggesting coordinated prefrontal control. In contrast, individuals with OCD showed increased connectivity between the RLPFC and MTG, indicating a more direct influence of RLPFC on posterior regions. Effective connectivity analyses further indicated that, across participants, the ACC/DLPFC and MTG function as central hubs of information flow, with task-related inputs entering the circuit via the TOJ, propagating through the MTG to the RLPFC, and subsequently modulating ACC/DLPFC and downstream regions. These findings suggest a shared underlying circuit architecture in OCD and healthy participants despite differences in functional coupling, particularly involving prefrontal cortical regions. Overall, differences arise at the level of functional coordination within a preserved circuit for abstract sequential processing in OCD, adding to current neurobiological models of OCD and suggesting a novel circuit that supports abstract sequencing.

## Introduction

Obsessive compulsive disorder (OCD) is characterized by complex symptomatology and likely supported by several networks of brain regions. OCD is a common psychiatric disorder, affecting 2-3% of the population, characterized by recurring, unwanted thoughts and accompanying actions or rituals (Stein et al., 2019). OCD symptoms can be complex and often manifest as dysfunctional performance of behavioral sequences. For example, repetitively washing hands, related to contamination concerns (Fontenelle et al., 2004), might disrupt a longer sequence of cleaning the dishes. Alternatively, a ritual such as excessively arranging books on a shelf may be conceptualized as dysfunctional over-engagement in behavioral sequencing. These behavioral patterns appear as disruptions in abstract sequences, a set of tasks defined by a rule rather than the identity of associated stimuli (Desrochers et al., 2022). While the role of cortico-striatal-thalamo-cortical (CSTC) loops are well-established in underlying OCD symptoms (Shephard et al., 2021), the potential contribution of circuitry specific to abstract sequencing is less well delineated.

Cortical regions and circuitry that support abstract sequencing have previously been established. A key node in a network of regions, the rostrolateral prefrontal cortex (RLPFC), was necessary for performance on an abstract sequence task in healthy participants (Desrochers et al., 2015). The blood-oxygen-level dependent (BOLD) response in this region monotonically increased from the start to the end of each sequence, a dynamic known as ramping activity. Ramping in the RLPFC is thought to potentially index sequential control or related processes based on findings from this and other studies (Desrochers et al., 2015, 2019; Doyle et al., 2025; McKim & Desrochers, 2022). Task-related ramping activity was observed across several frontal and parietal regions, as well, indicating a role for this network in abstract sequencing in healthy participants. The potential contribution of the RLPFC and other sequence-related circuitry to OCD symptomatology, however, remains unclear.

Previous work points to a potential network of cortical areas that are differentially activated in OCD compared to healthy participants during abstract sequencing. These regions may act as distinctive neural circuits that contribute to OCD symptoms involving sequential behavior. In one study, OCD participants exhibited several differences in neural activity dynamics during abstract sequencing (Doyle et al., 2026). Specifically, performing an abstract sequence task evoked increased ramping activity in OCD compared to healthy participants in a region spanning the rostral anterior cingulate cortex (rACC) and the superior frontal sulcus (SFS), the supplementary motor area (SMA), primary motor cortex and somatosensory cortex, parts of the middle (MTG), superior and posterior temporal gyrus, and the temporo-occipital junction (TOJ). Although these areas differentiate OCD from HCs during the task, it remains unclear whether they are functionally connected and the potential contribution of this circuit to OCD symptoms.

The shared ramping dynamic between the RLPFC and the ACC, SFS, SMA, MTG, and TOJ may reflect coordinated task-relevant cognitive processing within a circuit supporting abstract sequencing. These brain areas have been shown to be involved in processes related to abstract sequence performance. For example, neural oscillatory signals ramp in the DLPFC, a region within the SFS, during response inhibition (Khan et al., 2024) and in the ACC in preparation for switching tasks (Hyafil et al., 2009). These processes are likely critical for guiding behavior during complex sequences. Single neurons exhibit ramping firing rates in the SMA and motor cortex during motor sequences (Russo et al., 2020), suggesting that dynamics in these areas could help maintain the temporal structure of ordered actions in the context of abstract sequences as well. The posterior temporal gyrus is involved in monitoring the progression of information held in working memory (Davey et al., 2016), a process that may be necessary to keep track of abstract sequences as they unfold. The TOJ contributes to the integration of visual information (Rennig et al., 2024), supporting the incorporation of perceptual information needed to evaluate sequence progression.

Given their differential activation and potential roles in sequence processing, these cortical regions may work together in a circuit to execute sequences, differentially in OCD compared to healthy counterparts. This circuit may further include frontoparietal regions such as the RLPFC, as this region is necessary for and exhibits the same ramping dynamic during abstract sequencing (Desrochers et al., 2015) and is anatomically connected to the ACC and DLPFC (Haber et al., 2022). Understanding the connectivity between these regions and potential clinical differences will contextualize previously observed activation findings and elucidate the potential role for circuitry invoked in abstract sequencing in OCD.

Anatomical and functional connectivity studies support the existence of connections between cortical regions implicated in abstract sequential tasks. Prefrontal regions, for example, are connected to each other and other regions of the circuit. The RLPFC and DLPFC are anatomically connected to each other reciprocally (Haber et al., 2022), and both functionally project to the rostral ACC at rest (Jin et al., 2018). Further, both afferent and efferent anatomical connections arise from the DLPFC to the middle and superior temporal gyrus (Avalos-Alais et al., 2025). Functionally, the DLPFC and RLPFC are coactivated with the SMA during increasingly demanding cognitive tasks (Stiers & Goulas, 2018). Less direct anatomical connections exist between the SMA, MTG and TOJ regions of interest, but functionally, the middle MTG connects with the SMA while the posterior MTG projects to the middle occipital gyrus, which overlaps with the TOJ, at rest (Xu et al., 2015). Overall, connections between cortical areas previously implicated in abstract sequencing have been established through both anatomical and functional studies.

Although limited, existing functional connectivity studies in OCD during cognitive control tasks point to hyperconnectivity in cortical regions implicated in abstract sequencing. One study showed increased task-based connectivity in the frontoparietal and default mode networks, which include the SFS, RLPFC, and ACC respectively, across three cognitive flexibility tasks in OCD compared to healthy participants (Liu et al., 2023). Another study investigated coupling among several functional networks in OCD during a cognitive control task, finding increased connectivity in the SMA and the frontal operculum (Cocchi et al., 2011), a region located in the SFS/rACC cluster reported previously (Doyle et al., 2026). A third study showed increased effective connectivity between the DLPFC, a subregion of the SFS, and the ACC in OCD compared to HCs during cognitive control (Schlösser et al., 2010). Existing task-based connectivity studies on cognitive control in OCD suggest potential hyperconnectivity in networks that include cortical regions implicated in abstract sequencing.

Varying cognitive demands may influence the extent of differential connectivity in individuals with OCD. Specifically, regional connectivity may either remain stable across task conditions or vary with changing task demands. Connectivity that generalizes across the task suggests domain-general functioning (Duncan & Owen, 2000; Fedorenko et al., 2012, 2013), whereas regions that alter their activity and connectivity according to specific task conditions are considered domain-specific (e.g., Muhle-Karbe et al., 2014). Prefrontal cortical regions showing domain-general activation may serve as controllers of downstream regions within the sequence-processing circuit, consistent with prior models of cognitive control (Badre, 2008; Badre & D’Esposito, 2007; Miller, 2000; Miller & Cohen, 2001). In the present study, the RLPFC, SFS, and ACC may therefore function as domain-general controllers of the SMA, MTG, and TOJ, which appear sensitive to sequence complexity.

Multiple sources of evidence support the RLPFC, ACC and SFS potentially functioning as domain-general controllers. The RLPFC may assume a higher-level, modulatory role over other prefrontal cortical regions. With strong connections to other prefrontal regions, particularly the DLPFC, a region in the SFS, (Haber et al., 2022), the RLPFC is positioned to influence circuit dynamics indirectly. Further, RLPFC engagement has been linked to symptom severity in OCD (Doyle et al., 2026), suggesting functional relevance to the disorder. Prior work showing generalized ramping in the RLPFC across sequencing demands in healthy individuals further supports its role in abstract, higher-order control processes (Desrochers et al., 2015, 2019; McKim & Desrochers, 2022). Increased ramping in the ACC and DLPFC in individuals with OCD is observed throughout an abstract sequencing task, rather than being tied to specific task conditions (Doyle et al., 2026), suggesting domain-general involvement of these regions. This sustained engagement is consistent with their proposed roles in cortico-striato-thalamo-cortical (CSTC) loops, in which the DLPFC projects to downstream subcortical regions and the ACC is proposed to function as a hub for information flow and gating (McGovern & Sheth, 2017). More broadly, both regions have been shown to modulate activity in association cortices, including parietal and temporal areas (Jung et al., 2022). Together, these findings support the idea that the RLPFC, ACC and DLPFC may act as domain-general nodes to support a circuit for abstract sequencing differentially in OCD and healthy participants, with the RLPFC potentially modulating the ACC and DLPFC to influence the circuit indirectly.

The SMA, MTG, and TOJ are the cortical regions showing activation differences between groups that may exhibit domain-specific connectivity patterns varying by sequence complexity. Prior work shows that these regions are sensitive to changing task demands. For example, the SMA is increasingly activated under greater cognitive control requirements and with more complex motor demands (Al-Wasity et al., 2021; Dalla-Corte et al., 2015; Karabanov et al., 2023), while posterior MTG activation correlated with more demanding semantic retrieval (Davey et al., 2016). The temporo-parietal junction, which extends into the TOJ, is involved in shifting attention to unexpected stimuli (Indovina & Macaluso, 2004), often dependent on conditional demands. Consistent with these studies, our previous findings showed that the SMA, MTG, and TOJ exhibited greater ramping in OCD relative to healthy controls during simple compared to complex sequences (see Methods for details), with the opposite pattern observed in healthy controls (Doyle et al., 2026). Connectivity between these regions may similarly differ by sequence type across groups, with increased communication in healthy controls during more complex sequences to support higher task demands, and a reversed pattern in OCD that may contribute to symptomatology. The connectivity between SMA, MTG and TOJ may therefore be domain-specific, modulated by task complexity, alongside generalized top-down influence from ACC/DLPFC and RLPFC to coordinate the execution of abstract sequences.

Our hypotheses propose that the RLPFC, ACC/DLPFC, SMA, MTG and TOJ coordinate as a circuit to support abstract sequencing. Previous work supports hypotheses that these cortical regions are interconnected during abstract sequencing (**Figure 1**), while also suggesting that patterns of activation and connectivity may differ between groups. We hypothesize specifically that the superior frontal sulcus and rostral ACC (SFS/rACC) and the RLPFC will coordinate the sequencing circuit in a domain-general fashion differentially between groups. We further hypothesize the SMA, MTG and TOJ regions will be connected differentially between groups in different task conditions as a putative “abstract sequencing” circuit, based on ramping activation results in these areas from the same study. Based on studies showing contributions of bottom-up processes to cognitive control (Geng & Vossel, 2013; Leung et al., 2024; McMains & Kastner, 2011; Pessoa et al., 2002), visual task-related input may enter through the TOJ to modulate this circuit, influencing hyperconnectivity in OCD participants, similar to hyperconnectivity in functional networks observed during cognitive control tasks (Liu et al., 2023).

**Figure 1.**
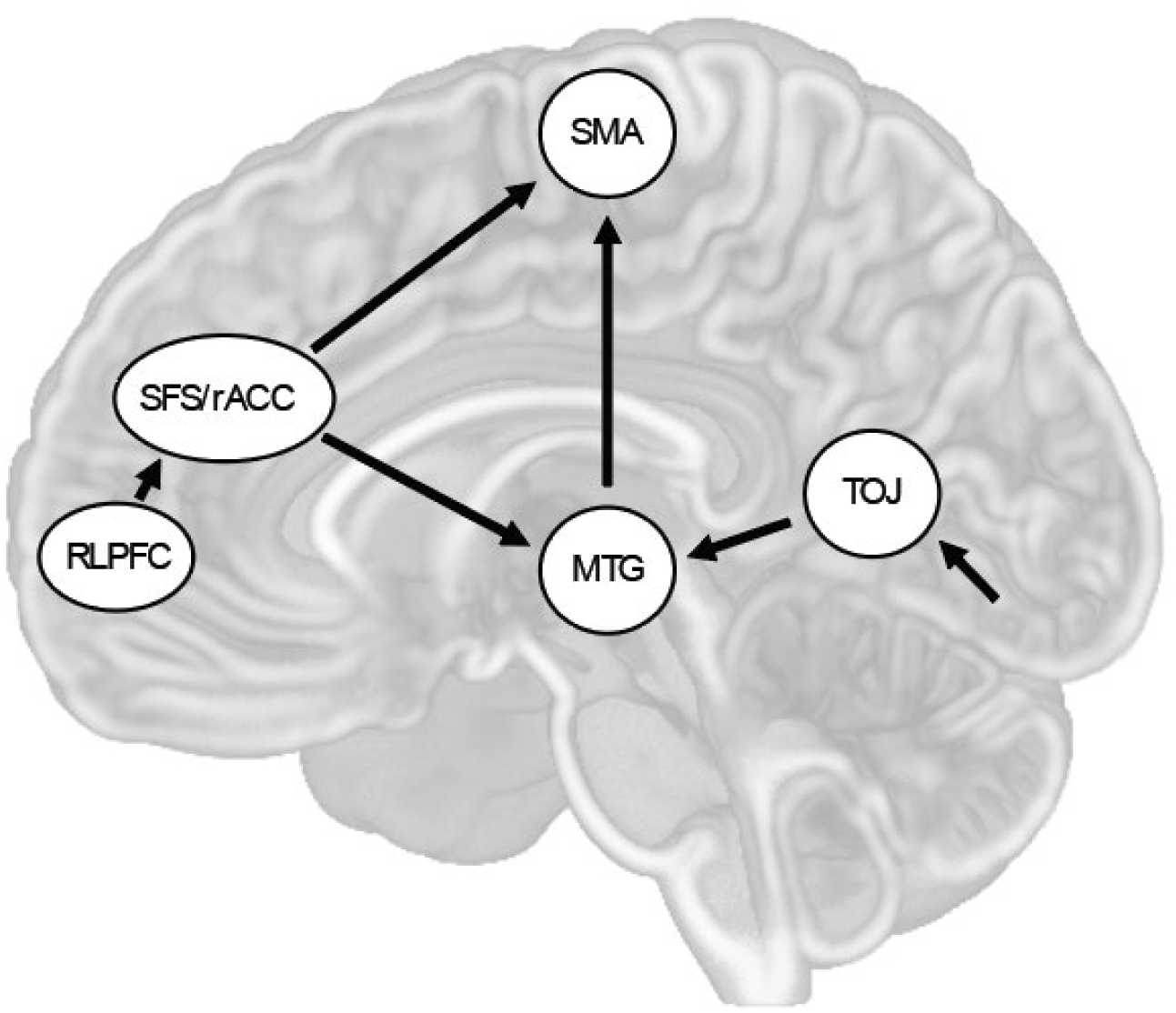
Hypothesis schematic of connectivity related to abstract sequencing. Based on literature and previous work in abstract sequencing, visual sequential input may enter the circuit through the temporo-occipital junction (TOJ), which is then passed to the middle temporal gyrus (MTG) and supplementary motor area (SMA) in a domain specific manner during different sequence conditions. Activation of this sequencing circuit may be controlled in a generalized way by the SFS/rACC region acting on the MTG or SMA, and further upstream by the rostrolateral prefrontal cortex (RLPFC).

To test these hypotheses, we used task-based seed-seed and seed-voxel functional connectivity (task-FC) and dynamic causal modeling (DCM) on functional magnetic resonance imaging (fMRI) data from a previous abstract sequencing study in healthy controls (HC) and OCD participants (Doyle et al., 2026). Overall, some regions connected differentially between groups, in a domain-general manner. The RLPFC was significantly positively connected to the MTG while the SFS/rACC was significantly negatively connected to the TOJ, both more in OCD than in HCs. The DCM revealed that across all participants, task conditions modulate the direction of connectivity such that input enters at the TOJ node, onto the MTG and rest of the circuit and potentially positions the RLPFC as more of a central hub within the circuit. Overall, these findings support the hypotheses of a specific sequence execution circuit that receives input from the TOJ and exhibits differential connectivity in OCD, particularly between the prefrontal regions SFS/rACC and RLPFC. Further, task input shifts connectivity to include RLPFC as a more involved node in the circuit. These findings elucidate differential cortical circuitry in individuals with OCD that is specific to abstract sequencing, in turn informing current neurobiological models conceptualizing networks implicated in this disorder.

## Methods

### Participants

Participant cohorts were from a previous study on abstract sequencing (Doyle et al., 2026). There were 25 participants in each group (HCs and OCD), and all 50 participants were included for FC and DCM analyses in the present study. Participants were originally recruited using fliers, word of mouth, and advertisements in the local community. OCD inclusion criteria largely matched that of HCs but additionally included a diagnosis of OCD, established via semi-structured clinical interview, and a score of 16 or higher on the Yale-Brown Obsessive Compulsive Scale (YBOCS) (Goodman, Price, Rasmussen, Mazure, Delgado, et al., 1989; Goodman, Price, Rasmussen, Mazure, Fleischmann, et al., 1989). Please refer to Doyle et al. (2026) for full study recruitment details and a full description of inclusion and exclusion criteria for each group. Final sample sizes and demographics of participants in this study are detailed in **Table 1**.

**Table 1.** Demographics for HC and OCD participant groups. Final sample size numbers reported for each group. Mean age (in years) and Y-BOCS scores were reported with +/- 1 SD (standard deviation). Note that no Y-BOCS scores were collected for HCs and thus were not reported. Total number of males and females were reported. Percentage identifying of a particular race was reported per group. Note that the total percentage adds to over 100% as participants were allowed to identify as more than one race. The total number identifying as Not Hispanic/Hispanic or Unknown identity were reported. Note for the HC group: Age and Sex information are reported for all 25 in the HC group (total number in this group). In the HC group, 2/25 participants completed this fMRI study prior to the clinical interview installment within the protocol, so additional demographic data (Race, and Ethnicity) were not collected and therefore not reported for these two participants. Table and description adapted from Table 1 in Doyle et al., (2026).

|  | HC | OCD |
| --- | --- | --- |
| **Final sample size (n)** | 25 | 25 |
| **Age** | 28.9 yrs. (+/- 10.7) | 25.8 yrs. (+/- 8.5) |
| **Sex** | 11 m (14 f) | 3 m (22 f) |
| **Race** | 87% White, 13% Asian, 1% Black | 84% White, <1% Asian, 12% Black |
| **Ethnicity** | 17(6) Not Hispanic (Hispanic) | 20(4)(1) Not Hispanic (Hispanic)(Unknown) |
| **Y-BOCS** | N/A | 22.28 (+/- 4.09) |

### Task Design and Procedure

#### Overview

Data were utilized from a previous fMRI study using an abstract sequence task (Doyle et al., 2026). In brief, participants were presented with a sequential rule at the start of each task block. This rule was used to make categorization decisions of color or shape on each trial. Each experiment consisted of 5 runs total, each with 4 blocks that contained 24-27 trials. Sequence rules were either ‘simple’ (‘AABB’ rule, containing one internal task switch) or ‘complex’ (‘ABBA’, containing two internal task switches). Each block consisted of only simple or only complex sequences within it and each run contained 2 simple sequence and 2 complex sequence blocks (**Figure 2A**).

**Figure 2.**
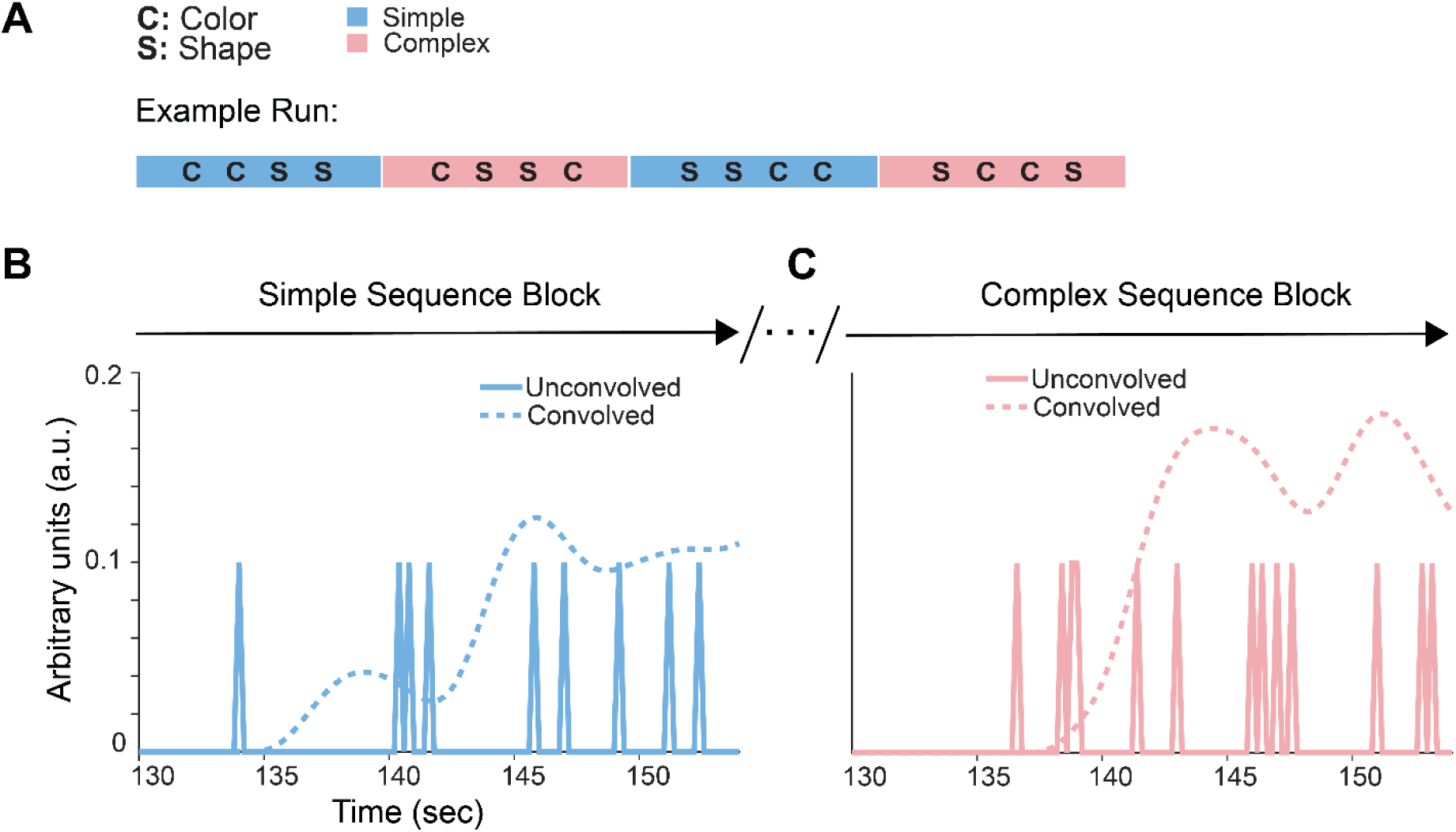
Illustration of abstract sequence task structure and task-FC design. **A.** Example run of complex and simple sequence blocks. Runs contained 2 blocks of complex sequences (rule ABBA: color, shape, shape, color or shape, color, color, shape) and 2 blocks of simple sequences (rule AABB: color, color, shape, shape or shape, shape, color, color). Simple sequences occurred only within simple sequence blocks (blue), and complex sequences were contained only within complex sequence blocks (pink). Conditions were defined by sequence block type. The “All Stimuli” condition was composed of all events from complex and simple sequence blocks. **B.** Example simple sequence block showing the first 2-3 sequences of the block and onset regressors used for task-FC design. Unconvolved onset regressors for each sequence position event were defined and input into CONN toolbox. These were then convolved with the canonical hemodynamic response function (HRF) to produce convolved regressors in the model. **C.** Same as in **B** but for complex sequences.

#### Data Acquisition

Data were acquired using a 3T PRISMA functional magnetic resonance imaging scanner with a 64-channel head coil. Functional data for two participants were collected using an echo planar imaging pulse sequence (repetition time, TR = 2.0 s; echo time, TE = 28 ms; flip angle 90°; 38 interleaved axial slices; 3.0 x 3.0 x 3.0 mm). The functional images for the remaining 48 participants were collected with different parameters intended to maximize signal to noise ratio (repetition time, TR = 1.53 s; echo time, TE = 33 ms; flip angle 62°; 60 interleaved axial slices; 2.4 x 2.4 x 2.4 mm). Anatomical data were collected with the same parameters across all participants. These images included a T1-MPRAGE (TR, 1900 ms; TE, 3.02 ms; flip angle, 9.0°; 160 sagittal slices; 1.0 x 1.0 x 1.0 mm) and a T1 in-plane scan (TR, 350 ms; TE 2.5 ms; flip angle, 70°; 38 transversal slices; 1.5 x 1.5 x 3.0 mm).

### Data Acquisition

#### Preprocessing

Task based data were preprocessed for the previous study using SPM12 in Matlab 2017b. Images were resampled for acquisition timing differences. They were then corrected for motion, and participants with more than one voxel of motion (3.0 mm for the first two participants, 2.4 mm for the remaining 48 participants) were excluded from further analysis. Data were then smoother using a Gaussian kernel. Please refer to Doyle et al., (2026) for more details of preprocessing data. Additional preprocessing for the present study was conducted in CONN toolbox 2022a for connectivity analyses (Whitfield-Gabrieli & Nieto-Castanon, 2012). This included denoising, a linear regression to remove confounding effects of white matter and cerebrospinal areas, and scrubbing, and a temporal band pass filter to isolate low frequence task related BOLD signal (range 0.008 - 0.1 Hz).

### ROI Definition

#### Functional connectivity

SMA, MTG and TOJ ROIs were determined by clusters of ramping activation in OCD compared to HCs in simple versus complex sequences (Doyle et al., 2026) (**Figure 3A-D**). The SFS/rACC activation cluster (Doyle et al., 2026) was split into three ROIs. These were determined by three peak activations in the cluster at the group level of analysis, each with each peak in a different anatomical region of the cluster to obtain a full representation of activation spread. Spheres were formed around those three peaks to produce 8 mm diameter ROIs (**Figure 3A-D**). These ROIs were called the ‘anterior dorsal SFS,’ (adSFS) in the rACC, the ‘anterior ventral SFS’ (avSFS) spanning the ACC and SFS regions, and the ‘posterior SFS,’ (pSFS) capturing more of the DLPFC and SMA (along the SFS).

**Figure 3.**
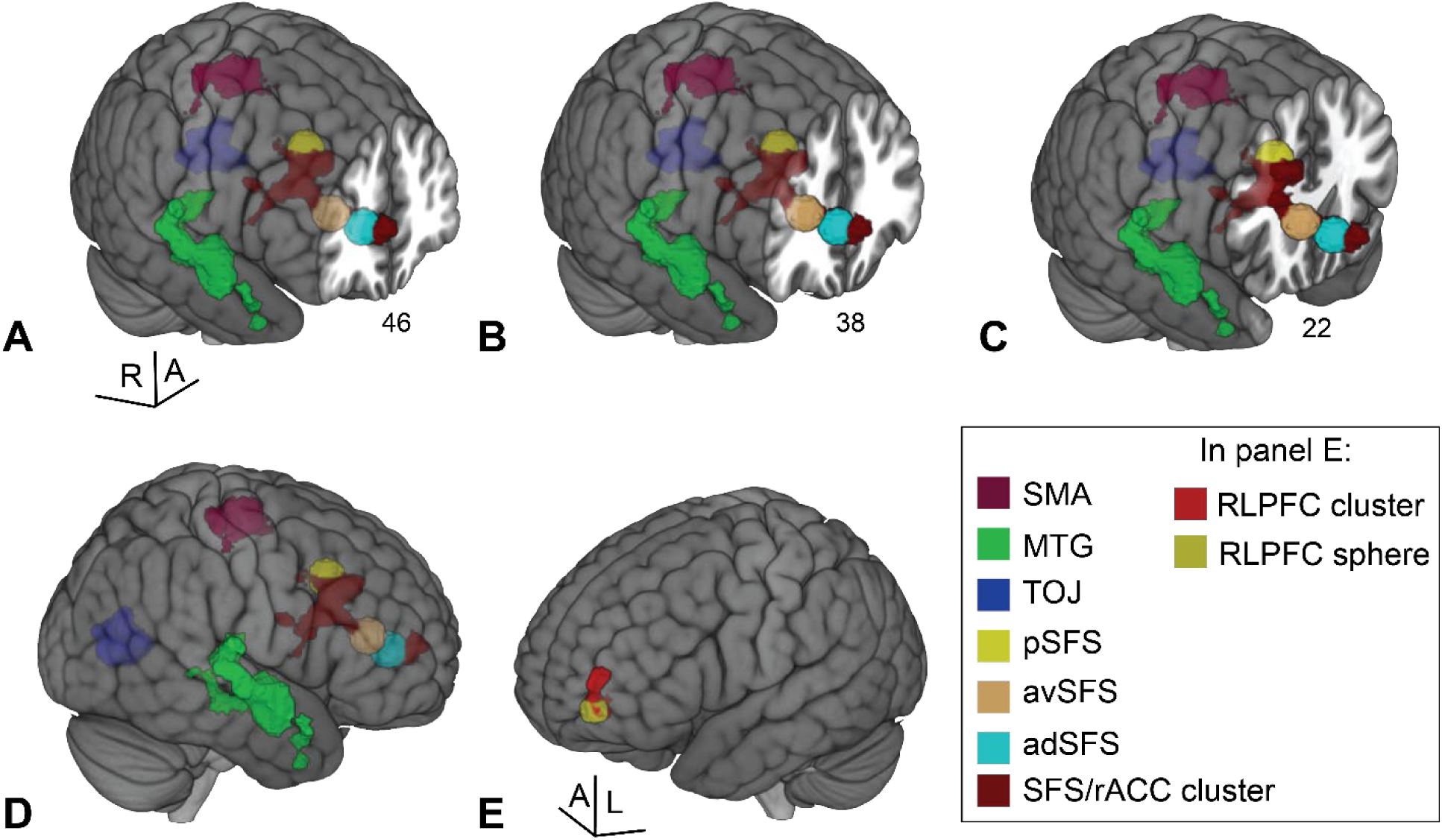
Location of ROIs for task-FC and DCM analyses. **A-E.** View of all spherical and cluster ROIs used for connectivity analyses on a reconstructed whole glass brain. The SFS/rACC cluster itself was not used as an ROI in connectivity analyses; rather, representative peaks of activation were used to form 8 mm spheres for task-FC and 6 mm spheres for effective connectivity (DCM) analyses. These anterior dorsal (adSFS), anterior ventral (avSFS), and posterior (pSFS) spheres are superimposed on the SFS/rACC activation cluster used to form them. SMA, MTG, and TOJ activation clusters were used in seed-seed and seed-voxel analyses. Note: spheres built for DCM analyses were based on the peak activation coordinates of clusters but were 6 mm in diameter (not shown). **A.** A coronal slice is shown to illustrate the center of the adSFS sphere. **B.** Same as (A) for avSFS sphere. **C.** Same as (A) for pSFS sphere. **D.** Right lateral view of the same ROIs in A, B, and C. **E.** Left lateral view of reconstructed whole glass brain showing only the 6 mm sphere RLPFC ROI (yellow) used for DCM analyses superimposed on RLPFC ROI cluster of activation (red) from Desrochers et al., (2015).

#### Dynamic causal modeling

ROIs were spheres defined from peak coordinates taken from the All Stimuli > Baseline and Simple > Complex OCD > HC (simple compared to complex sequences, greater in OCD compared to HC participants) ramping contrasts reported previously (Doyle et al., 2026). Specifically, for the SFS/rACC, peak coordinates matched those used in task-FC analyses. The RLPFC was defined as an additional ROI by forming a sphere around a peak coordinate from the All Ramp > Baseline contrast of a study investigating abstract sequencing in HCs (Desrochers et al., 2015) (**Figure 3E**). ROIs were defined as 6 mm spheres around individual participant activation peaks within 10 mm of the activation peak defined at the group level of analysis. Smaller spheres were chosen for DCM to better ensure nodes represent a single source of signal, an assumption of the model (Daunizeau et al., 2009). Following ROI definition, the first eigenvariate of the timeseries was extracted from each ROI per participant. The explained variance of each ROI was calculated per participant, and outliers outside of +/- 1 SD of mean were identified. In total, one outlier was identified, and this peak was manually adjusted to the coordinates of the group level peak to improve explained variance.

### Analysis

#### Functional connectivity

Seed-seed (ROI-ROI) and seed-voxel analyses were conducted in CONN toolbox, 2022a (Whitfield-Gabrieli & Nieto-Castanon, 2012). First level subject analyses included the following defined task conditions: “All Stimuli,” “Complex,” and “Simple.” In the “All Stimuli” condition, onsets and durations for events in complex and simple sequence blocks were used (trials in all blocks). In the “Simple” condition, onsets and duration for events in only the simple blocks were used (**Figure 2B**). In the “Complex” condition, onsets and durations for events in only the complex blocks were used (**Figure 2C**). These onset regressors were convolved with the canonical hemodynamic response function (HRF) by CONN toolbox (**Figure 2B,C**). Convolved regressors were then used as temporal weights to extract condition-specific BOLD signals across time to produce weighted timeseries. Condition-specific timeseries were then correlated between the seed and other voxels or seeds. Second level connectivity analyses included a covariate to determine the effect of group membership. Seed-seed analyses generated Fisher transformed correlation coefficient (r) values, plotted as heatmaps for illustration. Some seed-seed results were shown using boxplots (https://github.com/raacampbell/notBoxPlot). Significant results for seed-voxel analyses show Fisher transformed correlation coefficient (r) values that survive family-wise error (FWE) cluster correction (p < 0.05) at a height of p < 0.005.

#### Dynamic causal model (DCM

All DCM analyses were conducted using the Dynamic Causal Modeling built-in functions and tools in SPM12 (Friston et al., 2003). Broadly, a model was designed with nodes and connections between them hypothesized to best describe data. This model was fit to timeseries data by estimating parameters (parameter estimates, Ep) set in the model and posterior distributions of those parameters (the certainty of the estimates, posterior probabilities, Pp). The estimation of these parameters was conducted across subjects and between groups. The fully estimated model was then compared with alternative models to determine the best fitting model across subjects.

Single subject first level analyses were conducted by creating a general linear model for each participant, producing a BOLD timeseries for each voxel. The same onsets model used in task-FC analyses was used for DCM. In this model, the onset of each stimulus (i.e., image stimulus at each position of the sequence) is defined and set with a duration of zero seconds (Doyle et al., 2026). Each stimulus onset is labeled by its position number in the 4-item sequence and as part of the “Complex” or “Simple” sequence condition. In the DCM, the onsets for each condition were converted into input vectors in the model. In final analyses, parameter estimates were averaged across sequence positions within the Complex and Simple sequence conditions. Volumes of interest for each ROI were extracted for each participant (see details in ‘ROI definition’ section). The following matrices were specified in the model: fixed connectivity (A matrix), modulatory effects (B matrix), and direct task inputs to ROIs (C matrix). Bilateral connections were enabled, timeseries were used as data, and center input was enabled for all estimated models.

The DCM was then estimated at the first level for each participant using variational Bayes under Laplace approximation. After model estimation, DCM functions defined in SPM12 were used to determine how well the model explained individual subject data (explained variance). Following the guideline that explained variance should be approximately 10% per participant, this threshold was evaluated to ensure adequate model fit, as consistently low explained variance across participants could indicate that the model does not sufficiently capture the observed data overall (Friston et al., 2003; Stephan et al., 2010; Zeidman et al., 2019). Fifteen subjects with very low explained variance were identified (a model fit of less than 5%). To see if these subjects significantly impacted the model’s parameter estimates, a second level group analysis was conducted with all subjects included and a secondary analysis was conducted excluding the 15 low variance subjects. Model statistics were then compared between the two. Specifically, parameter estimates (Ep), the actual values of connections specified in the model, and posterior probabilities (Pp), a distribution indicating the certainty of the estimates (Pp of 0.95 and above used as a threshold to determine significance of Ep) were compared. Between the two models, most Ep differences (approximately 88%) were below 0.1, and approximately 13% of Pp were different. Therefore, most model evidence was similar whether including or excluding the 15 low variance participants. These numbers were used to justify including these subjects in the final sample.

The second level group analysis used parametric empirical Bayes (PEB), which treats first level DCM parameter estimates as noisy observations of a group-level generative model. In the fully estimated PEB, fixed connections were included between nodes based on task-FC results, and specific modulatory influences and driving effects on the TOJ node based on hypotheses. Fixed connections between nodes were assumed to be bi-directional. In the PEB step, a covariate was included to differentiate group membership (OCD and HC groups).

The final step of DCM was model comparison, to identify the best fitting model for the data. This step used Bayesian model reduction (BMR) and Bayesian model averaging (BMA) to compare the estimated second level PEB from the previous step with DCM template models. In the BMR step, all parameters in the model templates were set to zero, and Bayes factors were computed between nested models. If models were determined to be too similar, the model space was reduced. Seven model templates were included. Each template contained all fixed connections included in the estimated PEB (see **Figure 6**), and templates varied which node received driving effects from task input. The final template was a null model that included fixed connections but not modulatory or driving influences. The BMA step determined the presence of more than one dominant model. In this case, BMA averaged model parameters between the winning models to create one final best fitting model. This step does not apply if only one dominant model is observed among the templates. From the model comparison step, significant parameter estimates of the winning model across participants were identified and reported (**Table 5**; **Figure 7**).

## Results

Data were from a previous fMRI experiment in which participants executed abstract sequences (Doyle et al., 2026). In brief, participants were shown a four-item sequence (e.g., the words color, color, shape, shape) at the start of each block and used this rule to make categorical decisions (choosing the color or shape) of stimuli presented serially. Rules were categorized as simple (one internal task switch, ‘AABB’ rule) or complex (two internal task switches, ‘ABBA’ rule) sequences. Each block contained all simple or all complex sequences within it. Participants performed five runs, each containing four counterbalanced sequence blocks (two simple and two complex) (**Figure 2A**). In the present study, we used task based functional connectivity (task-FC) and dynamic causal modeling (DCM) to determine directed influences between cortical regions previously shown to be activated in OCD compared to HCs in this task (Doyle et al., 2026).

### Significant connectivity between nodes is present in HC and OCD groups

Because subsequent hypotheses relied on significant connectivity between groups, we first investigated FC in HC and OCD groups separately. We used seed-seed analyses to test correlations in the “All Stimuli” condition. This condition included all stimulus onsets of sequences from both “Complex” and “Simple” sequence blocks (**Figure 2**). The seeds of interest were chosen for their necessity in abstract sequencing (RLPFC) (Desrochers et al., 2015) and differential activity dynamics between OCD and HCs in the same task (anterior dorsal SFS, adSFS; anterior ventral SFS, avSFS; posterior SFS, pSFS; supplementary motor area, SMA; middle temporal gyrus, MTG; and temporal occipital junction, TOJ) (Doyle et al., 2026) (see **Figure 3** and Methods for details). In HCs, there were significant correlations among these regions. Significant positive connections were observed among the SFS clusters (anterior dorsal, anterior ventral, and posterior) and these clusters were also significantly connected to other more anterior and posterior regions: adSFS and pSFS with the RLPFC, adSFS with the SMA and MTG, and pSFS with the SMA and MTG (**Table 2**; **Figure 4A,B**). Additionally, there were significant correlations between the TOJ and the MTG, and between the SMA and the MTG. Significant negative correlations were observed between the TOJ and all the SFS clusters. In OCD participants, many correlations were consistent with those observed in HCs, with three differences: the RLPFC was not significantly connected to the pSFS, but was significantly positively correlated to the MTG, and the MTG was not significantly correlated to the TOJ (**Table 2**; **Figure 4C,D**). To summarize, across both groups, prefrontal regions exhibited positive connections between themselves and the SMA and MTG, and negative connections with the TOJ with differences in connectivity with RLPFC and MTG in OCD. We used these findings as the basis to test questions about group differences in connectivity across all nodes.

**Figure 4.**
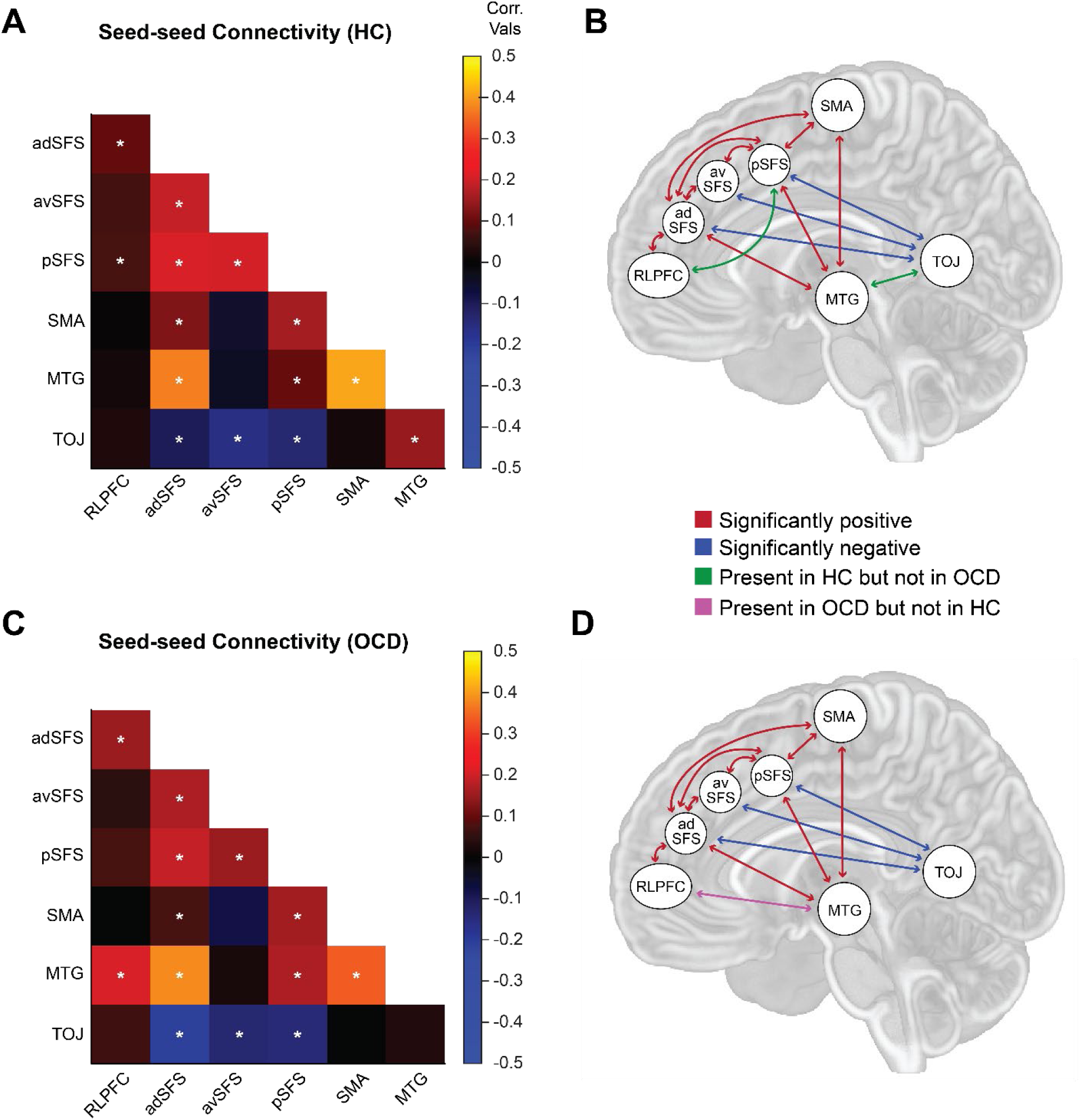
Seed-seed connectivity throughout the task (All Stimuli condition) for HC and OCD groups separately. **A.** Heatmap showing Fisher transformed correlation coefficients for each ROI-ROI pair in the HC group. Correlation values significantly different from zero (p < 0.05) are indicated with a white asterisk. **B.** Illustration of the significant ROI-ROI correlation values shown in (A). Green arrows indicate connections that are present in the HC but not the OCD group. **C.** Same as (A) for the OCD group. **D.** Illustration of the significant ROI-ROI correlation values shown in (C). Pink arrow indicates connection present in OCD but not in the HC group. **B, D**. Red arrows indicate signifcantly positive correlations, while blue arrows indicate significantly negative correlations.

**Table 2.**
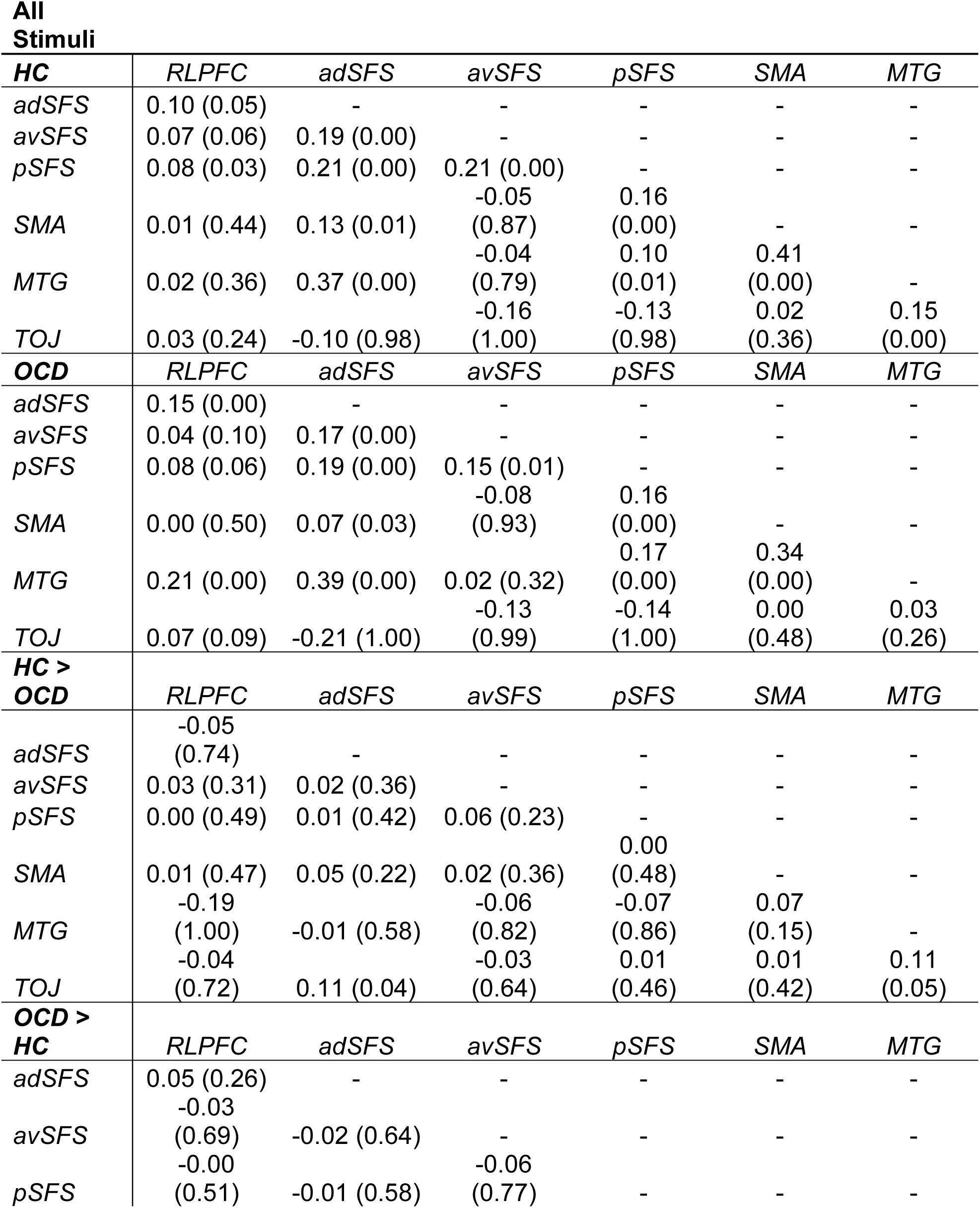
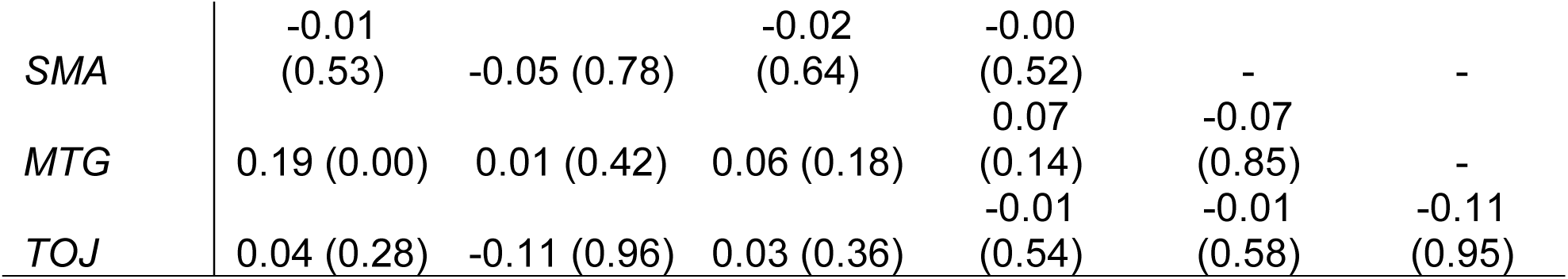
Seed-seed connectivity between all regions of interest in HCs and OCD separately and between groups, in the “All Stimuli” contrast. Correlation values (Fisher transformed correlation coefficients) for each pair are reported with associated significance (p values) in parentheses, following the order of presentation in the heatmaps in **Figure 4**.

### Differential connectivity between OCD and HCs occurs during abstract sequencing

We hypothesized differential connectivity between groups may occur between nodes either in a domain-general (i.e., consistently throughout the task) or domain-specific (in specific sequence conditions) manner. These hypotheses were largely based on how these regions were activated in a previous study. Using the same abstract sequence task, this study showed increased ramping in the SFS/rACC more in OCD than in HCs consistently throughout the task (Doyle et al., 2026). The SMA, MTG, and TOJ nodes, however, were differentially activated in simple and complex sequence conditions between groups. We therefore hypothesized the SFS/rACC and the RLPFC, another prefrontal controller region, would be connected to other nodes in the circuit throughout the whole task, while the SMA, MTG, and TOJ may be connected only during certain conditions and differently between groups. We tested these hypotheses first by examining group differences in connectivity throughout the task (in the “All stimuli” condition), and then between task conditions (comparing the “Complex” and the “Simple” sequence conditions).

We first tested hypotheses about domain-general connectivity differing between groups during the task. We used seed-seed analyses on all nodes across sequence types, implementing a covariate identifying group membership to detect group differences. Overall, we observed group differences in connections between both the RLPFC and SFS/rACC to other nodes during the sequence task. Connections with the RLPFC were positive in both groups, while connections to the SFS/rACC were negative. Specifically, the RLPFC was significantly more positively connected to the MTG in OCD compared to HCs (OCD > HC: z = 0.19, p = 0.003) (**Figure 5A,B**). In OCD participants, this connectivity was significantly greater than zero (t(24) = 4.39, p < 0.001), while in HCs it was not (t(24) = 0.37, p = 0.71) (**Figure 5C**), leading to this group difference. The adSFS was significantly more negatively connected to the TOJ in OCD compared to HCs (OCD > HC: z = −0.11, p = 0.04) (**Figure 5A,B**). Connectivity between these nodes was negative in both groups (HC: t(24) = −2.10, p = 0.05; OCD: t(24) = −5.11, p < 0.001) but significantly more so in OCD (**Figure 5D**). In addition to these hypothesized prefrontal regions, we observed a group difference between the MTG and TOJ in the “All Stimuli” condition. Specifically, connectivity between these nodes was significantly more positive in HCs than in individuals with OCD (OCD > HC: z = −0.11, p = 0.05), with HC connectivity significantly greater than zero (HC: t(24) = 3.23, p = 0.004; OCD: t(24) = 0.65, p = 0.52) (**Figure 5A,B,E**). Group differences between HCs and OCD (the inverse contrast) were the exact opposite of those shown in **Figure 5A**. Overall, we observed preferential involvement of prefrontal cortical regions with other nodes of the circuit between groups, potentially with RLPFC being more utilized in OCD while the SFS/rACC is more engaged for this purpose in HCs.

**Figure 5.**
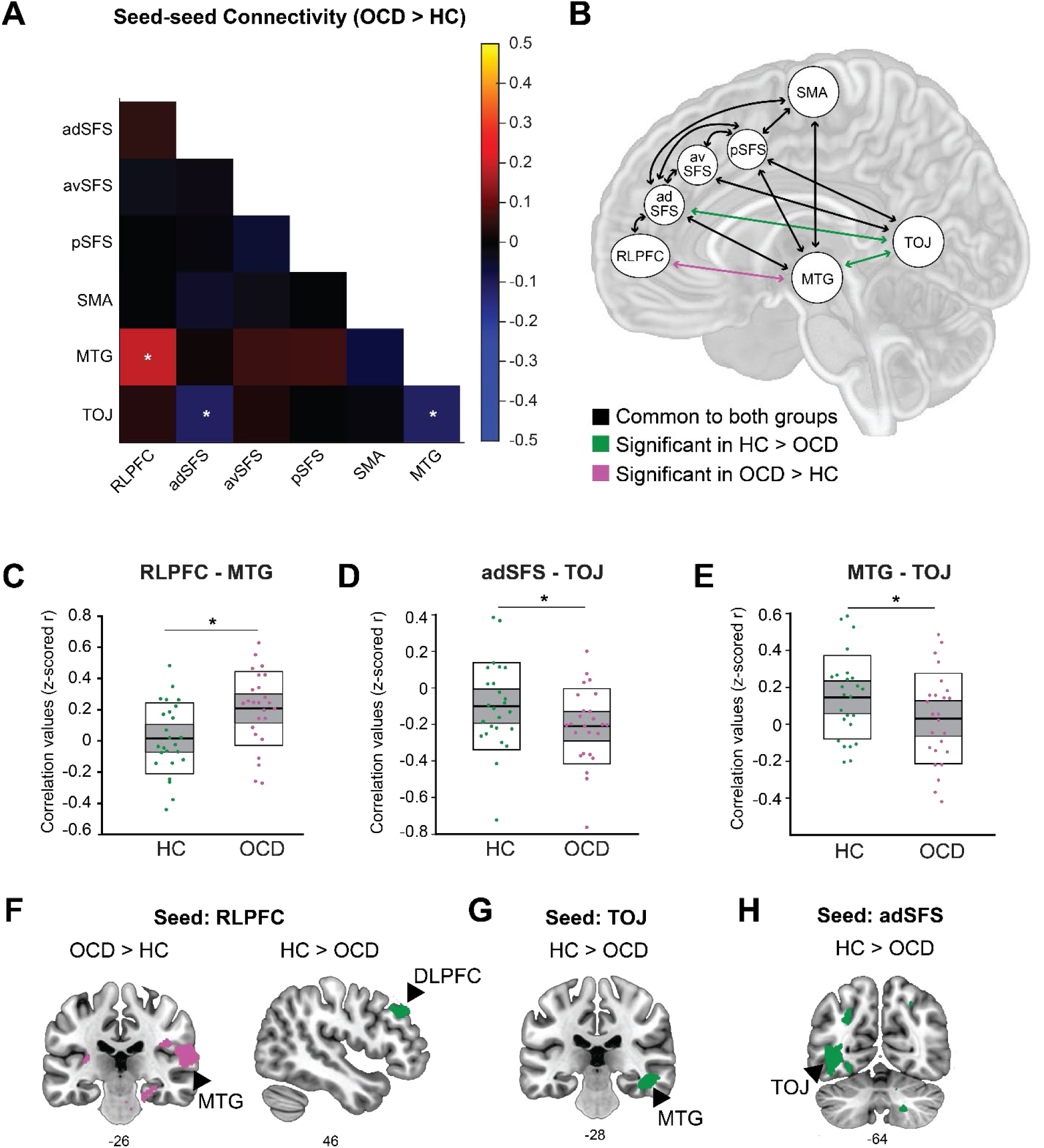
Seed-seed and seed-voxel analyses between participant groups during the sequence task (All Stimuli condition). **A.** Heatmap of beta values amongst all ROI-ROI analyses showing the difference between OCD and HC groups (OCD > HC). Significant differences (p < 0.05) between groups are indicated with a white asterisk. **B.** Illustration of functional connections common to both groups (black), significantly more in HC > OCD (green), and significantly more in OCD > HC (pink). **C-E.** Boxplots showing correlation values (Fisher transformed correlation coefficients) for each participant. Center solid black lines indicate data means, gray boxes indicate 95% confidence intervals, and outer white boxes with black borders indicate one standard deviation. HCs data shown in green, OCD shown in pink. Black asterisk indicates significant difference (p < 0.05) between groups. Three seed pairs shown: **C.** RLPFC-MTG. **D.** adSFS-TOJ. **E.** MTG-TOJ. **F.** Seed-voxel analysis in OCD > HC (left) and HC > OCD (right) using RLPFC as seed in the “All Stimuli” condition. Correlation values are plotted after FWE cluster correction, height p < 0.005, extent 241 (left) and 231 (right) voxels. **G.** Seed-voxel analysis in HC > OCD using TOJ as a seed in the “All stimuli” condition. Correlation values are plotted after FWE cluster correction, height p < 0.005, extent 270 voxels. **H.** Seed-voxel analysis in HC > OCD during the task using anterior dorsal SFS as seed, in the “All stimuli” condition. Connectivity correlation values are plotted at p < 0.05, uncorrected, for illustrative purposes only.

Findings from the seed-seed analyses are largely supported by whole-brain seed-voxel analyses. We conducted group-contrasted seed-voxel analyses using the adSFS, RLPFC, and TOJ as seed regions. We showed significant connectivity involving the RLPFC that paralleled seed-seed results and additional connections that were stronger in HCs than in OCD. Specifically, the RLPFC showed significant connectivity with a large region of the middle and superior temporal gyrus in OCD compared to HCs (FWE cluster corrected, p < 0.05, height p < 0.005, extent 241 voxels) (**Figure 5F**), replicating seed-seed findings, as well as other regions such as the parahippocampal gyrus and parts of the parietal cortex (**Table 3**). We additionally observed that the RLPFC significantly connected to a region of the dorsolateral prefrontal cortex (DLPFC) in HCs compared to OCD (FWE cluster corrected, p < 0.05, height p < 0.005, extent 231 voxels) (**Figure 5F**; **Table 3**). We additionally replicated seed-seed results of differential connectivity between the MTG and the TOJ. The MTG significantly connected to the TOJ during the task more in HCs than in individuals with OCD in seed-voxel analyses (FWE cluster corrected, p < 0.05, height p < 0.005, extent 270 voxels) (**Figure 5G**; **Table 3**). We did not observe significant seed-voxel connectivity from the adSFS to TOJ that was greater in HCs compared to OCD as we did in seed-seed analyses. However, at lower thresholds (p < 0.05, uncorrected), this region does connect to the TOJ in HC > OCD, which we show for illustrative purposes (**Figure 5H**). Overall, these findings support seed-seed results between groups and point to differential roles for the RLFPC in influencing the circuit between HCs and OCD participants (see Discussion).

**Table 3.** Seed-voxel results from “All Stimuli” contrast using RLPFC and TOJ as seeds, between HC and OCD groups. Clusters reliable at p < 0.05 FWE cluster corrected. Extent p < 0.005 for all contrasts. All clusters listed with distance between significant clusters > 12 mm. Coordinates are the center of mass in MNI.

| Contrast Location | BA | Extent<br>(voxels) | x | y | z | Peak t-val. |
| --- | --- | --- | --- | --- | --- | --- |
| **All Stimuli, RLPFC seed** |  |  |  |  |  |  |
| **OCD > HC** |  |  |  |  |  |  |
| Central operculum | 48 | 1958 | 64 | -4 | 12 | 4.67 |
| Planum temporale<br>(superior temporal<br>gyrus) | 22 |  | 64 | -26 | 14 | 4.56 |
| Precentral gyrus | 4 |  | 44 | -8 | 36 | 4.3 |
| Supramarginal gyrus | 48 |  | 62 | -38 | 22 | 3.78 |
| Precentral gyrus | 48 |  | 42 | -8 | 22 | 3.76 |
| Planum temporale<br>(superior temporal<br>gyrus) | 41 |  | 48 | -36 | 18 | 3.64 |
| Parietal operculum | 48 |  | 42 | -26 | 26 | 3.32 |
| Planum polare | 38 |  | 52 | 4 | -4 | 3.16 |
| Central operculum | 48 |  | 50 | -4 | 10 | 2.96 |
| Postcentral gyrus | 6 |  | 32 | -16 | 40 | 2.81 |
| Brain stem | NA | 241 | 20 | -22 | -26 | 3.87 |
| Parahippocampal gyrus | 30 |  | 26 | -30 | -16 | 3.81 |
| Planum temporale | 42 | 242 | -56 | -42 | 16 | 3.84 |
| Parietal operculum | 48 |  | -36 | -34 | 16 | 3.39 |
| **HC > OCD** |  |  |  |  |  |  |
| Dorsolateral prefrontal<br>cortex (DLPFC) | 45 | 231 | 48 | 36 | 36 | 4.59 |
| Middle frontal gyrus | 8 |  | 36 | 26 | 56 | 3.59 |
| Middle frontal gyrus | 45 |  | 56 | 32 | 26 | 2.96 |
| **All stimuli, TOJ seed** |  |  |  |  |  |  |
| **HC > OCD** |  |  |  |  |  |  |
| Medial orbital frontal<br>cortex | 11 | 270 | 18 | 52 | -16 | 4.7 |
| Anterior orbital gyrus | 11 |  | 32 | 58 | -18 | 3.58 |
| Medial orbital frontal<br>cortex | 11 |  | 8 | 58 | -22 | 3.33 |
| Hippocampus/inferior<br>temporal gyrus | 20 | 281 | 42 | -24 | -16 | 5.54 |
| Posterior cingulate<br>gyrus | 30 | 468 | 14 | -46 | 18 | 3.79 |
| Cuneus | 18 |  | 18 | -74 | 32 | 3.68 |
| Precuneus | 18 |  | 14 | -46 | 36 | 3.56 |
| Precuneus | 18 |  | 22 | -54 | 24 | 3.44 |
| Cerebellum | NA | 646 | -26 | -90 | -22 | 3.76 |
| Cerebellum | NA |  | -14 | -80 | -34 | 3.7 |
| Cerebellum | NA |  | -34 | -76 | -40 | 3.25 |

### Differences in connectivity do not depend on specific sequence task conditions

Based on previous univariate findings that the SMA, MTG and TOJ showed differences in activation between groups specific to sequence type (simple vs. complex), we hypothesized that these areas would be significantly connected in a domain specific manner, i.e. depending on sequence type. Our previous study found these regions ramped in activity significantly more in OCD than in HCs in simple compared to complex sequences (Doyle et al., 2026). It is possible that ramping indexes sequencing-related cognitive processes that support performance as task demand increases (Davey et al., 2016; Rennig et al., 2024; Russo et al., 2020). Therefore, we hypothesized that these regions would significantly project to each other differently between groups and task conditions. Using seed-seed analyses with a focus on these regions, we did not observe significant differences in connectivity between groups and task conditions, between these nodes or any other regions of interest (**Table 4**). Overall, group differences in connectivity occurred in a domain-general manner throughout the task (**Figure 5**) and were not influenced by specific task conditions.

**Table 4.**
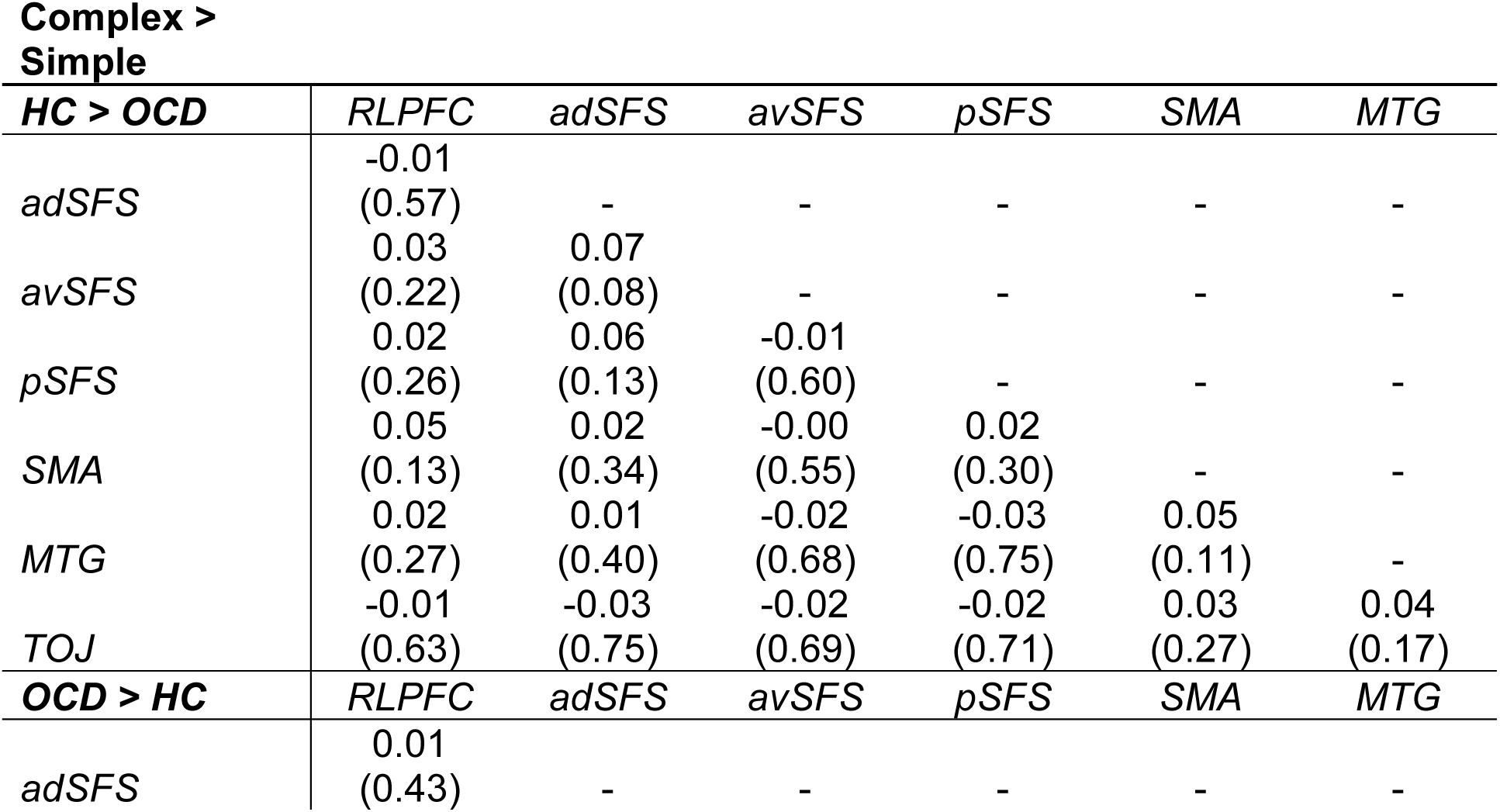
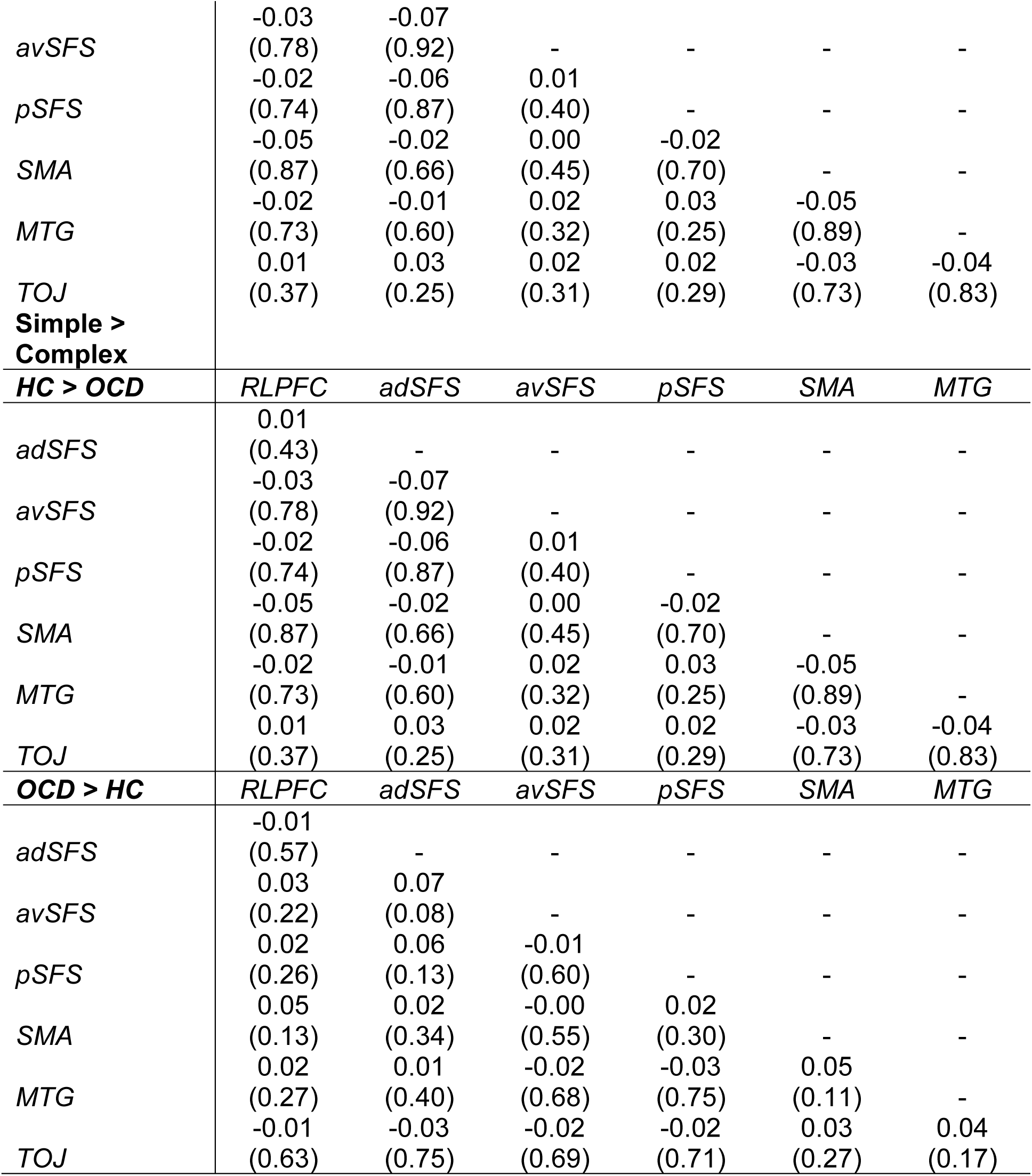
Seed-seed connectivity between all regions, between groups and between conditions contrasts. Correlation values (z scored r values) for each seed-seed pair are reported with associated significance (p values) in parentheses, between groups (HC > OCD and OCD > HC) and between sequence type conditions (Complex > Simple and Simple > Complex). No seed-seed pair correlations were found to be significant.

### Dynamic causal modeling supports proposed circuitry implicated in abstract sequencing

We next investigated the directionality of established connectivity between nodes using dynamic causal modeling (DCM). While DCM as a method has wide-ranging applications, we used the model as a tool to answer specific questions. First, we tested if connectivity patterns observed between groups were associated with task modulation of specific nodes. Second, we tested if driving effects from task-related stimuli acted on specific nodes in the circuit. These hypotheses were based on previous univariate results suggesting that specific task conditions may modulate connectivity between nodes of interest differentially in OCD and HCs (Doyle et al., 2026). To answer these questions, we designed a model with fixed connections between nodes based on FC seed-seed findings (**Figure 6**). We tested if effects from the sequence task (i.e., Complex or Simple conditions) modulated specific connections. Based on FC results, we hypothesized that MTG-TOJ and RLPFC-pSFS connections would be more strongly modulated by the task in HCs compared to OCD, while RLPFC-MTG would be more strongly modulated in OCD. We additionally tested the hypothesis that driving effects from the task would enter the circuit at the TOJ node, based on literature suggesting bottom-up processing of visual stimuli contributes to higher cognitive processes (e.g., McMains & Kastner, 2011; Pessoa et al., 2002). We tested this model compared to alternatives that modeled driving input at every other node, and a null model that did not include modulatory or driving effects (see Methods).

**Figure 6.**
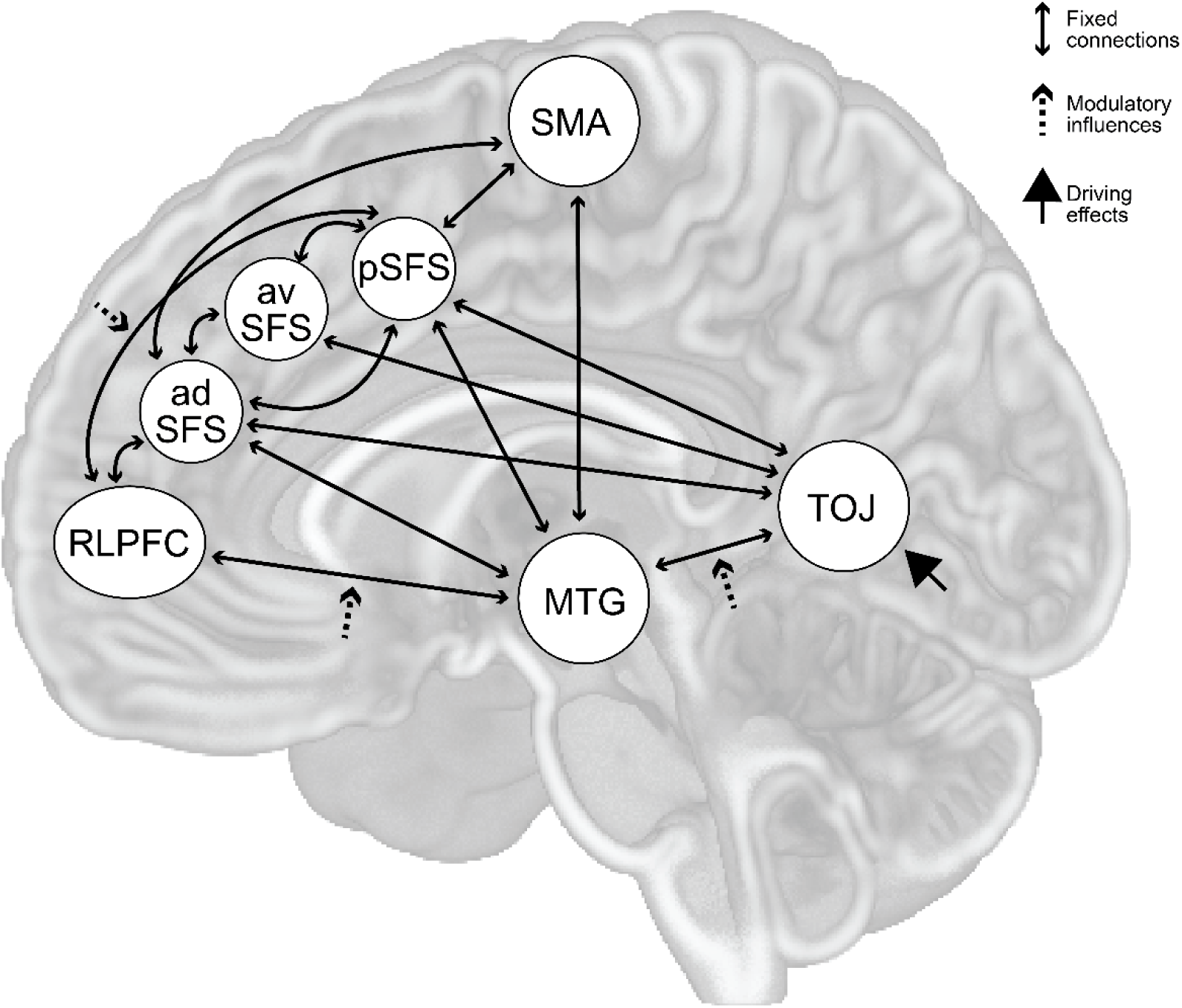
Illustration of hypothesized DCM connections. Nodes and fixed connections (double ended arrows) between them drawn from seed-seed FC results. Modulatory connections (influences from the task, dashed arrows) were allowed on connections that were shown to be different between groups in the seed-voxel FC analyses. These connections were between the RLPFC and adSFS, the RLPFC and MTG, and the TOJ and MTG. Hypothesis model included driving effects (solid head arrow) from the task entering the model at the TOJ node.

The DCM revealed a model that best fit data across all participants that largely aligned with our hypotheses. Specifically, Model 1 (**Figure 6**), best fit data across participants, indicating preference for driving input entering via the hypothesized TOJ region rather than alternative nodes. Model comparison identified redundancies and reduced the model space to three comparable models: the full hypothesis model illustrated in **Figure 6** (Model 1), a model with modulatory connections but no driving effects (Model 2), and a null model that did not include modulatory influences or driving effects (Model 3). The model comparison revealed Model 1 best fit data across all participants, independent of group membership (100% probability).

Parameter estimates and posterior probabilities for fixed connections, modulatory influences, and driving effects are reported in **Table 5**, and their effects are illustrated in **Figure 7**. Fixed connections (solid arrows) showed that on average during the task, most nodes were significantly connected to each other. The majority of connections were excitatory, with some negative directed connectivity between the adSFS and SMA, and from the avSFS to the TOJ and from the SMA to the MTG. Fixed connections also showed that the SFS/rACC (particularly the pSFS node) and the MTG may have been situated as central hubs within the circuit, mainly directing excitatory influences onto other nodes, while receiving some input as well.

**Figure 7.**
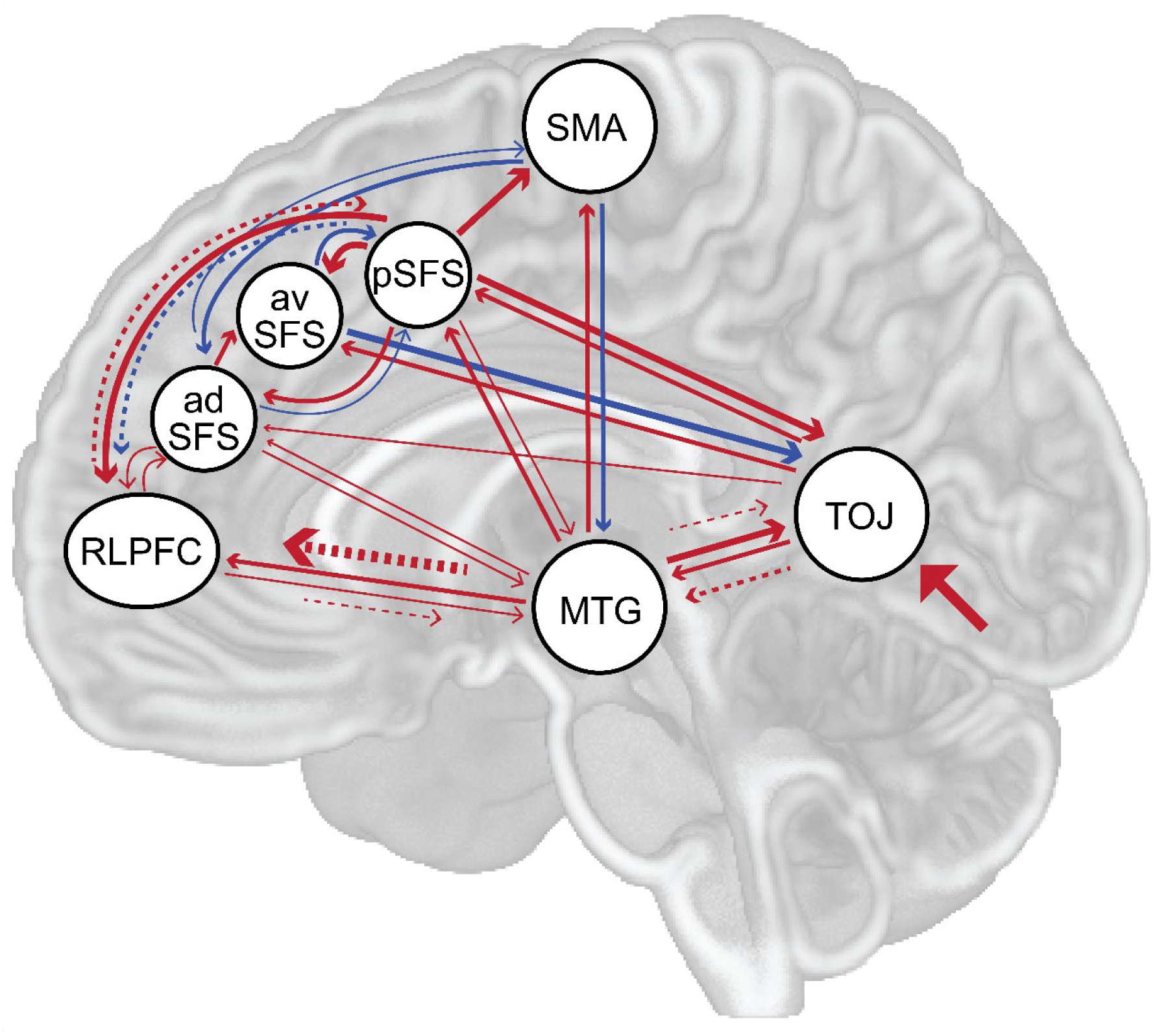
Illustration of effective connectivity produced by significant parameter estimates for the best fitting DCM model across all participants. All fixed connections except three (avSFS ◊ adSFS, pSFS ◊ SMA, and adSFS → TOJ) had significant parameter estimates. All modulatory connections and driving effects in this model had significant parameter estimates for at least some task conditions (see **Table 5** for details). Valence of connectivity is indicated by line color (red = excitatory, blue = inhibitory), and line thickness illustrates strength of a connection/influence on a node. Parameter estimates were averaged across sequence conditions for readability and because no FC differences were observed between sequence types. Please refer to **Table 5** for all parameter estimates in the model, broken down by sequence type when relevant.

**Table 5.**
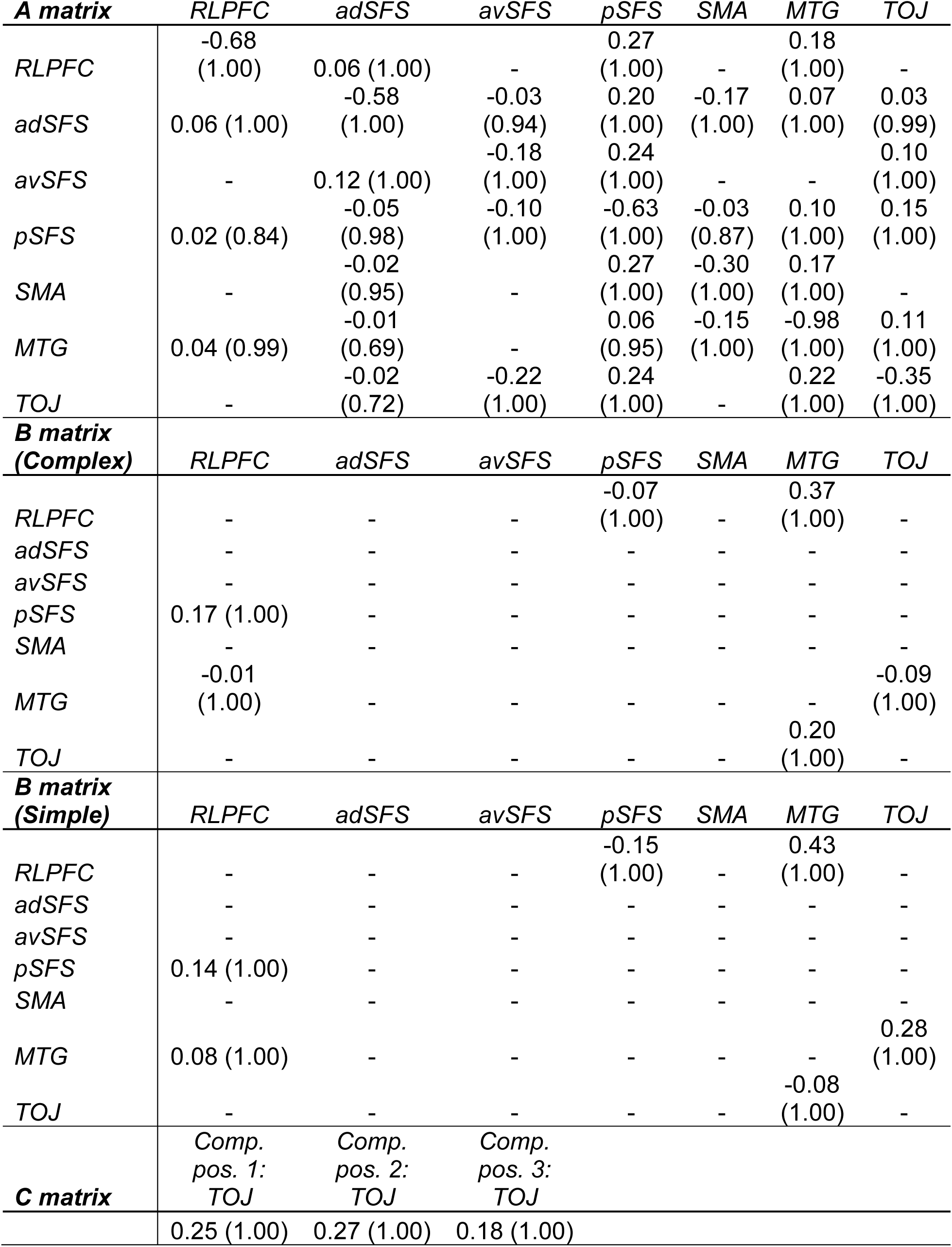
Parameter estimates and posterior probabilities for all fixed connections (A matrix), modulatory (B matrix) and significant driving (C matrix) influences in the winning DCM, across participants. Direction of connectivity determined by reading column ROI ◊ row ROI in A and B matrices. B matrices are displayed for Complex and Simple sequence conditions, separately. C matrix displays conditions with significant parameter estimates. These included sequence positions 1, 2, and 3 in the Complex condition, when the driving effect was on the TOJ node. Non-significant C matrix estimates were not listed because their associated posterior probabilities were not available. Parameter estimates are listed with associated posterior probabilities in parentheses for all three matrices.

Modulatory task influences, in part, potentially position the RLPFC as more of a central node within the circuit. Task influences on connections (illustrated in **Figure 7** with dotted arrows) directed excitatory influences from the TOJ to the MTG and onto the RLPFC, which in turn was positively connected to the pSFS. These modulatory task influences therefore emphasized the role of the RLPFC as a key node for engaging the broader circuit, as well as the importance of connectivity directed from the TOJ to upstream circuit nodes. Driving effects also served to position the RLPFC as a hub of information flow in the circuit. In conjunction with modulatory task influences, driving task effects (illustrated in **Figure 7** with filled in arrow) enter the circuit at the TOJ node, which directed connectivity towards the MTG and to the RLPFC. Overall, the DCM supported a priori predictions about task influences on the direction of connectivity between nodes across all participants, suggesting that the task engaged information flow through nodes starting at the TOJ and included the RLPFC as a central hub (see Discussion).

We additionally tested group differences in effective connectivity using the same models. DCM model comparison reported Model 3, the null model, as best describing group differences (100% certainty). We therefore did not find that modulatory influences or driving effects on these nodes sufficiently explained connectivity differences between groups (see Discussion).

## Discussion

We investigated the relationship between cortical regions previously implicated in abstract task sequences using functional and effective connectivity methods. Specifically, we hypothesized that these task active regions (the RLPFC, SFS/rACC, SMA, MTG, and TOJ) would be differentially connected in OCD compared to HC participants, some in a domain-general and some in a domain-specific manner. We further hypothesized these cortical regions would comprise a circuit with task input entering at the TOJ node and prefrontal areas influencing downstream regions. We broadly found support for our hypotheses, observing support for a coordinated circuit of cortical regions during abstract sequencing with differences in connectivity between groups. Specifically, the adSFS and MTG were more connected to the TOJ in HCs than in OCD, while the RLPFC was more connected to the MTG in OCD compared to HCs. We further observed that task influences modulated the circuit such that input entered at the TOJ whereby modulatory influences increased connectivity to the RLPFC, which, in turn, increased the SFS/rACC connectivity in the circuit. These findings may indicate that prefrontal regions shape downstream processing within an abstract sequencing circuit, with differential roles of the SFS/rACC and RLPFC across groups, and task-driven modulation shifting the balance of prefrontal cortical influence to position the RLPFC as a central hub. More broadly, this work frames neural circuitry implicated in OCD as dynamically changing according to task engagement, suggesting that prefrontal hub regions such as the SFS/rACC and the RLPFC may be important to consider in identifying future treatment targets.

Connectivity findings broadly align with literature on functional network imbalances in OCD. Our observation that most cortical regions in each group separately had significant connections was supported by anatomical and task-FC evidence (Avalos-Alais et al., 2025; Haber et al., 2022; Jin et al., 2018; Stiers & Goulas, 2018; Xu et al., 2015). Connectivity between groups revealed hyperconnectivity between certain nodes in OCD, suggesting our findings generally fit previously established evidence regarding imbalances in functional network communication (Cocchi et al., 2011; Liu et al., 2023; Schlösser et al., 2010). In OCD, the adSFS, a node that overlaps with the rostral ACC, showed stronger negative connectivity with the TOJ, while the RLPFC had stronger coupling with the MTG compared to HC participants. Since the ACC and TOJ are associated with the default mode network while the SFS and RLPFC contribute to the frontoparietal network, it is possible group differences may reflect inter-network communication dysfunction in OCD. For example, disrupted salience network function alters connectivity with the frontoparietal network in OCD (Yu et al., 2024), which may relate to MTG and RLFPC connectivity differences observed during the task. Further, the default mode network, which includes the ACC and parts of the TOJ, is differentially connected to frontoparietal regions during cognitive control in OCD (Liu et al., 2023), as well. Overall, our connectivity findings are supported by previously established anatomical and functional connections between nodes, and group effects may reflect broader inter-network communication differences in OCD.

Across the task, significant task-FC suggests prefrontal nodes maintain different roles in the circuit in both groups, with an imbalance in their recruitment in OCD. In both groups, the adSFS was negatively connected to the TOJ while the RLPFC was significantly positively connected to the MTG. Connectivity between prefrontal regions and other circuit nodes in opposing directions suggests there may be different but complimentary roles for these regions to balance the rest of the circuit. Prior work has dissociated roles for the DLPFC, a region the SFS nodes overlap with, and RLPFC during decision-making. One study, for example, showed that while these regions work together, the DLPFC distinctly accumulates evidence while the RLPFC contributes higher-level cognitive information during decision-making (Shekhar & Rahnev, 2018). It is plausible that these prefrontal regions have similar coordinated, yet distinct, roles in making choices during abstract sequencing. In OCD, this balance is more extreme, with these participants exhibiting more negative adSFS and more positive RLPFC connectivity with downstream nodes. Group differences therefore support an imbalance in recruiting the SFS/rACC and RLPFC during abstract sequencing in OCD, aligning with frontoparietal dysfunction observed previously (Liu et al., 2023). Our prior work also supports this interpretation as we observed increased ramping in the SFS/rACC in OCD compared to HCs during this task (Doyle et al., 2026), which may compensate for its stronger negative connectivity with other circuit nodes. Together, connectivity findings support differential roles for the rACC/SFS and RLPFC to balance the rest of the circuit, with a potential imbalance observed in participants with OCD.

Effective connectivity findings suggest task influences shift the RLPFC to become a more central hub in the circuit across all participants. Findings from DCM showed that fixed connections between nodes place the SFS (particularly pSFS) and the MTG as central nodes in the circuit. However, adding input from the abstract sequence task in the model appears to shift information flow through the RLPFC which, in turn, acts on the SFS/rACC. These findings suggest a potential mechanism for how sequential input maintains a balance between the SFS/rACC and RLPFC to influence the rest of the circuit. The recruitment of the RLPFC during the task is supported by evidence that this cortical region is needed for abstract categorization choices and more complex tasks (Desrochers et al., 2015), such as the abstract sequencing task. Overall, effective connectivity findings suggest that task influences cause the shift in balance between the SFS/rACC and RLPFC nodes in coordinating with the rest of the circuit.

Both functional and effective connectivity results highlight the role of prefrontal regions as controllers of downstream circuit nodes. Previous work has implicated the DLFPC and RLPFC as hierarchical control regions, with the DLPFC involved in handling immediate task demands (Braver et al., 2009; Duncan & Owen, 2000; Kerns et al., 2004; Miller & Cohen, 2001), while the RLPFC manages higher-level abstract control demands (Badre, 2008; Badre & D’Esposito, 2007; Desrochers et al., 2015, 2016, 2019). Studies have causally implicated these regions in higher-order cognition. One, for example, showed that directly stimulating the DLPFC enhanced adaptive cognitive control in humans (Gbadeyan et al., 2016), while our previous work showed disruption of the RLPFC inhibits abstract sequencing performance (Desrochers et al., 2015). Our connectivity findings suggest the SFS/rACC and RLPFC nodes are similarly positioned as controllers, balancing positive and negative connections with other circuit nodes and directing information to downstream regions.

Effective connectivity findings show that task input enters the circuit to promote bottom-up information flow. The dynamic causal modeling results showed that across participants, driving effects from the task enter the circuit at the TOJ node, directing information flow to upstream nodes. In turn, prefrontal nodes (the SFS/rACC and RLPFC nodes) communicate back with downstream regions with a top-down influence. Both bottom-up and top-down influences are observed during cognitive control (Buschman & Miller, 2007; Corbetta et al., 2008; Katsuki & Constantinidis, 2014), with bottom-up information often entering through sensory regions to orient attention, including through the temporo-parietal junction (Geng & Vossel, 2013; McMains & Kastner, 2011) while top-down control from prefrontal regions contributes to goal-directed processes (Badre, 2008; Koechlin et al., 2003; McMains & Kastner, 2011). Our findings align with literature in supporting task input entering and driving the circuit in a bottom-up fashion.

The architecture of the abstract sequencing circuit appears to be preserved across task conditions, rather than showing strong domain-specific connectivity. Although we hypothesized that connectivity among the SMA, MTG, and TOJ would vary as a function of task demands, the observed group differences instead reflected a domain-general pattern, with stronger connectivity between the MTG and TOJ in healthy participants relative to those with OCD across task conditions. Prior findings (Doyle et al., 2026) demonstrated that local ramping activity in these regions varies across groups and task conditions, which, together with the present results, suggests that while regional processing is sensitive to task demands, the broader circuit organization remains stable across the task. This interpretation aligns with literature on the multiple demand network, a proposed collection of frontal and parietal regions invoked in a domain-general manner across cognitive tasks (Duncan, 2010). Studies investigating this network have found that nodes within it are sensitive to changing task demands and code specific task-related features or contexts, while the network itself exhibits preserved functional coupling across cognitive tasks (e.g., Stiers & Goulas, 2018; Woolgar & Zopf, 2017). In accordance with this literature, it is plausible that condition-specific ramping activity reflects modulation of local computational demands within individual nodes, while group differences in connectivity primarily arise from how prefrontal regions, particularly the SFS/rACC and RLPFC, are differentially recruited to coordinate the preserved circuit across task conditions, rather than from condition-dependent changes in the overall circuit structure.

Limitations to this study largely arise from expected variation in the clinical sample and indirect measures of causality. The variation in OCD symptom dimensions, severity, medication status and comorbidities inherent amongst clinical populations may mask observations in both functional and effective connectivity. Beyond the natural variability within clinical populations, task-FC and DCM provide estimates of statistical and directed interactions between nodes but offer limited insight into the underlying mechanisms that give rise to these interactions. This limitation arises in part from the indirect nature of the BOLD signal, which reflects a mixture of neuronal and vascular processes (Drew, 2019; Guilbert et al., 2022; Howarth et al., 2021) rather than a direct measure of neural activity. Accordingly, DCM inferences depend on assumptions about neural dynamics and neurovascular coupling, which may constrain the extent to which estimated parameters uniquely reflect underlying neuronal mechanisms of interactions between circuit nodes. An additional limitation of the DCM approach is that it cannot distinguish between a true absence of differences between clinical and healthy groups and a failure to detect group differences due to limited statistical power. Despite these limitations, the present findings provide evidence for a neural circuit supporting abstract sequencing, with connectivity alterations in OCD participants.

Future work may produce meaningful translational benefits by refining our understanding of the neural circuitry underlying abstract sequencing and its relevance in OCD. Currently, approved neuromodulation treatment for OCD includes transcranial magnetic stimulation (TMS) targeting the ACC (Carmi et al., 2019), and several studies have investigated its use on the DLPFC (Dehghani-Arani et al., 2024; Elbeh et al., 2016; Lusicic et al., 2018), yet the mechanisms supporting its efficacy remain incompletely understood. The present findings delineate a cortical circuit that incorporates the ACC and DLPFC, offering a more complete circuit-level account of how this treatment produces an effect. Future studies could evaluate the stimulation of additional circuit hubs, such as the RLPFC and MTG, particularly in individuals whose symptoms reflect dysfunctional sequential processing. Overall, this work proposes a plausible cortical circuit supporting abstract sequencing, provides evidence for altered recruitment of prefrontal regions in OCD, and advances circuit-based models of the disorder while identifying potential new targets for neuromodulatory interventions.

## Data Availability

Imaging data are available upon request. Analysis code is available upon request.

## Author Contributions

Hannah Hyde: Data Curation, Conceptualization, Investigation, Methodology, Formal Analysis, Writing – Original Draft, Writing – Reviewing and Editing, Visualization. Sarah Garnaat: Conceptualization (primary study), Methodology, Investigation & Training in Clinical Assessments, Clinical Resources, Writing – Reviewing and Editing, Supervision of Clinical Activities, and Funding Acquisition. Nicole McLaughlin: Methodology, Investigation & Training in Clinical Assessments, Clinical Resources, Writing – Reviewing and Editing, and Supervision of Clinical Activities. Theresa M. Desrochers: Conceptualization, Methodology, Validation, Writing – Original Draft, Writing – Reviewing and Editing, Supervision, Funding Acquisition.

## Acknowledgements

We thank members of the Desrochers Lab for their feedback on the manuscript and Dr. Derek Nee for methodological consultation on dynamic causal modeling.

